# Ranked tree shapes, non-random extinctions and the loss of phylogenetic diversity

**DOI:** 10.1101/224295

**Authors:** Odile Maliet, Fanny Gascuel, Amaury Lambert

## Abstract

Phylogenetic diversity (PD) is a measure of the evolutionary legacy of a group of species, which can be used to define conservation priorities. It has been shown that an important loss of species diversity can sometimes lead to a much less important loss of PD, depending on the topology of the species tree and on the distribution of its branch lengths. However, the rate of decrease of PD strongly depends on the relative depths of the nodes in the tree and on the order in which species become extinct. We introduce a new, sampling-consistent, three-parameter model generating random trees with covarying topology, clade relative depths and clade relative extinction risks. This model can be seen as an extension to Aldous’ one parameter splitting model *β*, which controls for tree balance) with two additional parameters: a new parameter *α* quantifying the correlation between the richness of a clade and its relative depth, and a parameter *η* quantifying the correlation between the richness of a clade and its frequency (relative abundance or range), taken herein as a proxy for its overall extinction risk. We show on simulated phylogenies that loss of PD depends on the combined effect of all three parameters, *β, α* and *η*. In particular, PD may decrease as fast as species diversity when high extinction risks are clustered within small, old clades, corresponding to a parameter range that we term the ‘thin ice zone’ (*β* < –1 or *α* < 0; *η* > 1). Besides, when high extinction risks are clustered within large clades, the loss of PD can be higher in trees that are more balanced (*β* > 0), in contrast to the predictions of earlier studies based on simpler models. We propose a Monte-Carlo algorithm, tested on simulated data, to infer all three parameters. Applying it to a real dataset comprising 120 bird clades (class Aves) with known range sizes, we show that parameter estimates precisely fall close to close to a ‘thin ice zone’: the combination of their ranking tree shape and non-random extinctions risks makes them prone to a sudden collapse of PD.

## Introduction

As it becomes increasingly clear that human activities are causing a major extinction crisis (Leakey and Lewin 1995; Glavin 2007; Wake and Vredenburg 2008; Barnosky et al. 2011), several theoretical studies have aimed at characterizing how the evolutionary legacy of parts of the Tree of Life, and hence also the genetic diversity able to drive future evolution, will decrease in the face of forthcoming extinctions. This evolutionary component of biodiversity can be measured by the phylogenetic diversity (PD), defined as the sum of the branch lengths of the phylogeny spanned by a given set of taxa (Faith 1992). This metric is increasingly being used to measure biodiversity and to identify conservation strategies (Veron et al. 2015).

Nee and May (1997) were the first to formally investigate the expected loss of PD in the face of species extinctions, by simulating species trees using the Kingman coalescent. They found that 80% of the phylogenetic diversity can be conserved even when 95% of species are lost. Further studies showed that the loss of PD is in fact much higher when trees are generated through other models of species diversification, such as the Yule or the birth-death models (Mooers et al. 2012; Lambert and Steel 2013). These models indeed generate longer pendant edges (*i.e*., branches that lead to the tips), hence lower phylogenetic redundancy, than in the standard Kingman coalescent (used by Nee and May 1997). However, Nee and May (1997) also showed that phylogenetic diversity is very sensitive to the shape of the species tree (also called its ‘topology’), with extremely unbalanced trees (‘comb trees’) losing much more phylogenetic diversity than balanced trees (‘bush trees’), due to a lack of phylogenetic redundancy (i.e., the presence of recently diverged sister species). Overall, these results highlighted the sensitivity of the loss of phylogenetic diversity in response to species extinctions to both edge lengths and tree shape.

In this line, we also expect the correlation between the species richness of clades and their relative ages to have a significant impact on the loss of PD (‘clade’ standing here for any subtree within the full phylogeny). Here the age of a clade, also called ‘stem age’, denotes the depth (measured from the present) of its root node (i.e., the node where this clade is tied to the rest of the tree). Under random extinction, since smaller clades are more likely to become extinct first, the consequence of their total extinction on PD will depend on the lengths of pendant edges in these clades compared to those in larger clades. The effect of such correlation on the loss of PD has not yet been explored, but should be particularly important in unbalanced phylogenetic trees (exhibiting large variation in the species richness of clades), which dominate empirical data (*e.g*., Guyer and Slowinski 1991; Heard 1992; Guyer and Slowinski 1993; Slowinski and Guyer 1993; Mooers 1995; Purvis 1996; Mooers and Heard 1997; Blum and François 2006).

Besides, the loss of PD was shown to be influenced by the distribution of extinction risks within species trees. Several studies showed that accounting for realistic scenarios of species extinctions (considering that species with higher extinction risk–as per the IUCN Red List status–are more likely to go extinct first) predicts proportionately higher losses in PD than scenarios with random extinction risks (*e.g*. and review, Purvis et al. 2000a; von Euler 2001; Purvis 2008; Veron et al. 2015). Extinctions may for example be clustered within certain clades (Bennett and Owens 1997; McKinney 1997; Russell et al. 1998; Purvis et al. 2000a; Baillie et al. 2004; Bielby et al. 2006; Fritz and Purvis 2010), correlated to the age of clades (von Euler 2001; Johnson et al. 2002; Redding and Mooers 2006), or to the species richness of clades (Russell et al. 1998; Hughes 1999; Purvis et al. 2000a; Schwartz and Simberloff 2001; von Euler 2001; Johnson et al. 2002; Lozano and Schwartz 2005, assuming in some studies a correlation between rarity and extinction risks). In contrast, theoretical analyses of predictions based on model trees (Nee and May 1997; Mooers et al. 2012; Lambert and Steel 2013) have all been based so far on the field of bullets model, which considers equal extinction probabilities across species (Raup et al. 1973; Van Valen 1976; Nee and May 1997; Vazquez and Gittleman 1998). One can assume extinction events are independent but not identically distributed across species, as considered in the generalized field of bullets model (Faller et al. 2008). In an exchangeable phylogenetic model in which extinction probabilities are themselves random and independent with the same distribution, this would not affect the overall loss of phylogenetic diversity (as both models are stochastically equivalent, Lambert and Steel 2013). However, as stated by Faller et al. (2008), it is essential to explore models that weaken the strong assumption in the (generalized) field of bullets models that extinction events are randomly and independently distributed among the tips of phylogenetic trees.

Here, we hence investigate how the loss of PD is influenced by the two abovementioned factors: (i) the ranked shape of the species tree, considering notably correlations between clade richness and clade depth, and (ii) non-random extinctions, considering notably correlations between clade richness and extinction risks within the clade. Here, ‘ranked shape’ refers to the shape of the tree combined with the additional knowledge of relative depths–the order in which nodes appear in the tree, but to the exclusion of the actual divergence times(*e.g*., Lambert et al. 2017).

We introduce a three-parameter model generating random ranked tree shapes endowed with random numbers summing to one at the tips, interpreted as relative abundances (or geographic ranges) of contemporary species. This model can be seen as an extension to Aldous’ *β*-splitting model (Aldous 1996, 2001) with two additional parameters: a parameter *α* quantifying the correlation between clade richness and clade relative depth (*i.e*., the rank in time of its root node; termed ‘correlation clade size-depth’ hereafter), and another parameter *η* quantifying the correlation between clade richness and its frequency (i.e., its relative abundance compared to that of all extant species in the phylogeny; termed ‘correlation clade size-frequency’ hereafter). When *β* = 0 and *α* = 1, the ranked shape of the tree is the same as the ranked shape of a standard coalescent tree or of a Yule tree stopped at a fixed time (see Proposition 1 in Appendix 1). We further assume that contemporary extinctions occur sequentially by increasing order of abundance, which roughly reduces to the field of bullets model when *η* = 1 (see Proposition 2 in Appendix 1).

We explore the rate of decrease of PD as species sequentially become extinct, based on simulated data under variation in all three parameters over a significant range of their possible values. Interestingly, the joint variation of the parameter *η* with the ranked shape of species trees (set by parameters *β* and *α*) affects the clustering of extinction risks and the relationship between extinction risks and clade depth (determined by the similarity or dissimilarity of the direction of deviations of *α* and *η* from 1). Therefore, considering simultaneous variation in *β, α* and *η* allows us to explore the effects on the loss of PD of the different patterns of non-random extinctions observed in empirical data. We therefore provide general predictions on the sensitivity of the evolutionary legacy of clades to extinction, as a function of three simple statistics summarizing tree balance, ranked tree shape and the distribution of extinction risks across clades.

Besides, we then propose a Monte-Carlo inference algorithm enabling maximum likelihood estimation of the parameters *β, α* and *η* from real datasets. When tested against simulated data, this algorithm performs reasonably well over a wide range of parameter values for phylogenies with 50 tips or more. The estimates of parameters (beta, alpha, eta) on a real dataset of bird family phylogenies and their range size distributions finally reveal empirical patterns clustered within a given parameter zone which make these clades particularly prone to strong loss of phylogenetic diversity.

## Methods

### Modeling ranked tree shapes

The first version of the model we present allows one to generate random ranked tree shapes, that is tree shapes endowed with the additional knowledge of node ranks. Usually, one can generate random ranked tree shapes by time-continuous branching processes stopped at some fixed or random time, where particles are endowed with a heritable trait influencing birth and death rates. In these models, it is generally not possible to characterize the distribution of the tree shape (for an exception, see Sainudiin and Véber 2016) or to relate it to known distributions whenever it does not have the shape of the Yule tree (i.e., the tree generated by a pure-birth process). Also, since the same trait is usually responsible for both the tree shape and the order of nodes, it is impossible to disentangle the roles of either of these characteristics on the behavior of the tree in the face of current extinctions. Last, these models do not fulfill a very important property called sampling consistency (usually considered in combination with exchangeability, i.e., ecological equivalence between species). This property ensures that one can equivalently draw a random tree with *n* tips from the distribution or draw a tree with *n* + 1 tips and then remove one tip at random.

The model we propose here has two parameters: *β* ∈ (–2, +∞) determines the balance of the tree, similarly as in Aldous’ *β*-splitting model (Aldous 1996, 2001), and *α* ∈ (–∞, +∞) sets the correlation between species richnesses of clades and their relative depths (Fig. 2).

**Figure 1:**
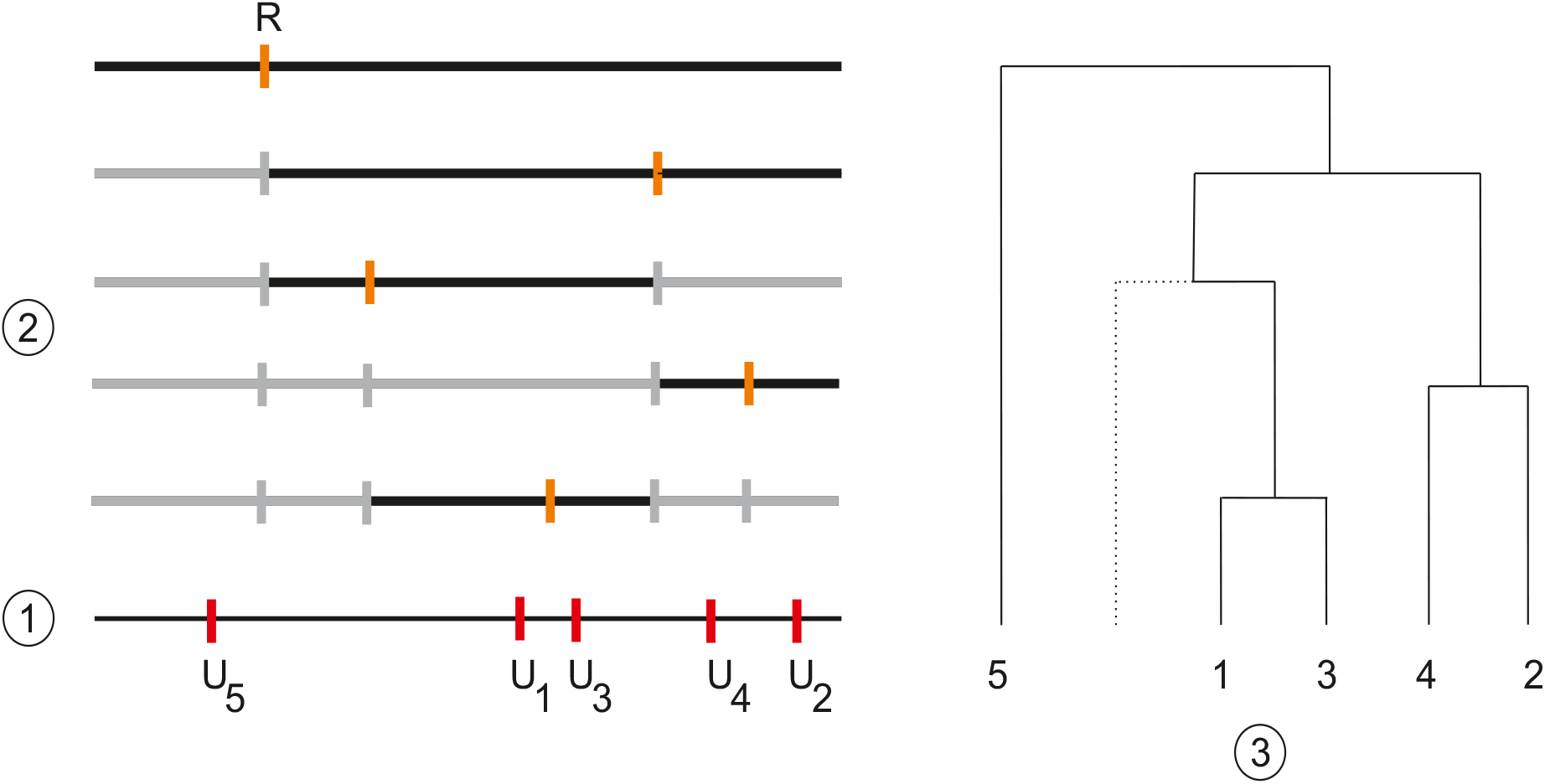
Illustration of the model generating ranked tree shapes. Construction of the ranked shape of a tree containing *N* = 5 species. (1) Five random marks (*U_i_*)_*i*∈{1,…,5}_ are drawn uniformly in the interval [0, 1] (red marks). (2) At each time step (time flowing downwards), we randomly select one interval *X*, with each interval *X_j_* having a weight |*X_j_*|^*α*^ (in black). Then, we draw a random variable *R* in a Beta distribution with parameters (*β* + 1, *β* + 1), and split the selected interval *X* into two subintervals, *X_left_* of size *R*|*X*| and *X_right_* of size (1 – *R*)|*X*| (orange mark). (3) Repeating this process over time until all intervals *X_j_* contain only one mark leads a tree with a ranked shape. Dotted branches correspond to unsampled subtrees (i.e. there is no mark in the corresponding interval).

**Figure 2:**
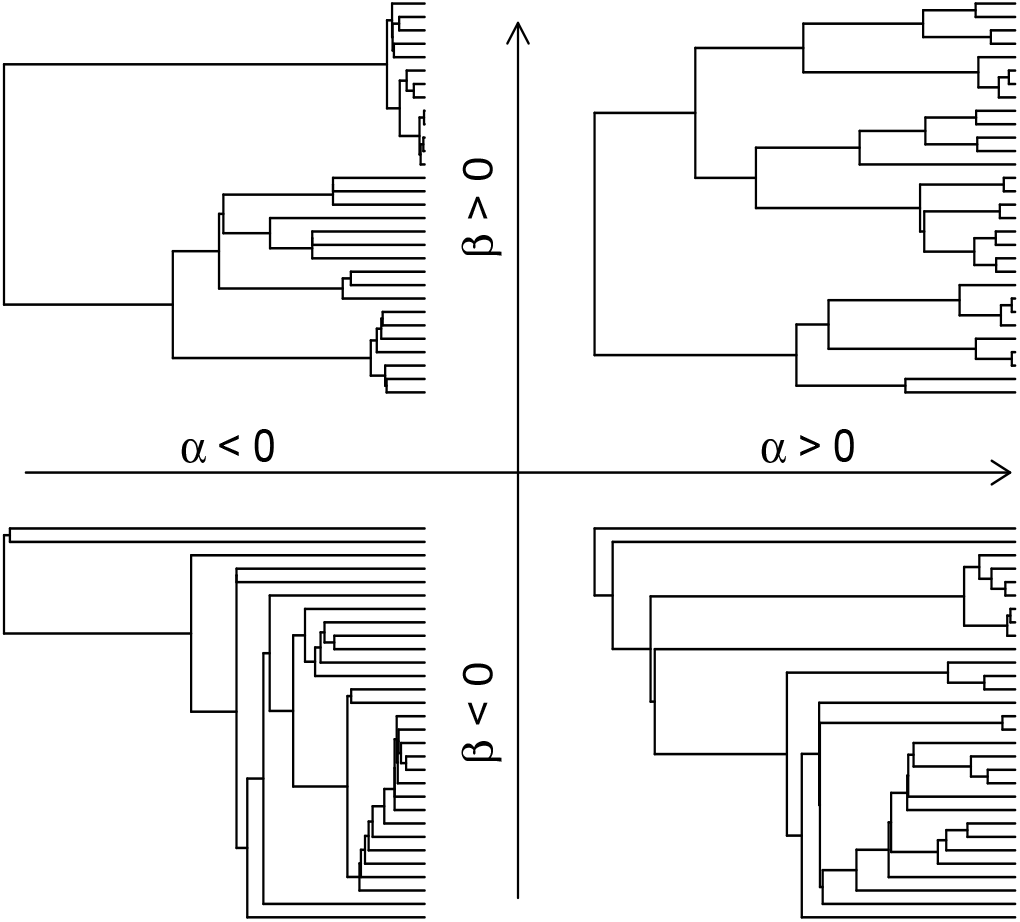
Phylogenetic trees simulated for different values of *β* (tree balance) and *α* (correlation clade size-age). Node depths are set as in a Yule pure-birth process. Parameter values: *β* = –1.5 (bottom) or 10 (top), *α* = –10 (left) or 10 (right), number of species *N* = 30, *ϵ* = 0.001.

The construction of a tree according to this model is done by following the steps indicated hereunder (illustrated on Fig. 1). We start with n uniform, independent random variables (*U_i_*)_*i*∈{1,…, *n*}_ in the interval [0, 1]. Each mark *U_i_* is associated to the tip species labelled i in the phylogeny. The procedure consists in sequentially partitioning [0, 1] into a finite subdivision thanks to random variables independent of the marks (*U_i_*)_*i*∈{1,…,*n*}_, until all marks are in distinct components of the partition. At each step, the new point added to the subdivision corresponds to a split event in the tree. In the beginning, there is only one component in the partition (the interval [0, 1] itself).

1. Each interval *X* of the partition containing at least two marks among the (*U_i_*)_*i*∈{1,…, *n*}_ is given a weight equal to |*X*|^*α*^, where |*X*| denotes the width of *X*. Then one of these intervals is selected with a probability proportional to its weight.
2. Draw a random variable *R* in a Beta distribution with parameters (*β* + 1, *β* + 1). The selected interval *X* of width |*X*| is then split into two disjoint subintervals, *X_1eft_* and *X_right_*, with widths |*X_left_*| = *R* |*X*| and |*X_right_*| = (1 – *R*) |*X*|. Each subinterval contains a distinct subset of the marks. The marks in the subinterval *X_left_* determine the tips in the left subtree of the phylogeny, and the marks in the subinterval *X_right_* determine the tips in the right subtree. This step is performed even if one subinterval contains no mark among the (*U_i_*)_*i*∈{1,…,*n*}_, which corresponds to a subtree with no sampled species. The order in which the splitting subintervals are selected sets the order of branching events (*i.e*., nodes) in the tree.
3. If no interval contains more than one mark, the process is stopped. Otherwise, go to Step 1.

We can relate the tree shape in this model to well-known distributions. Because *α* has no impact on the way we refine the subdivision, the tree shape generated with our model coincides with the tree shape with parameter *β* in Aldous’ *β*-splitting model (Aldous 1996, 2001). For small values of *β*, the intervals are often split close to an edge, and the resulting tree is imbalanced, converging to the perfectly imbalanced ‘comb’ tree as *β* → – 2. On the contrary, for large values of *β*, the intervals are often split close to the middle, and the resulting tree is balanced. We stress that unlike most models, *α* can be tuned independently of *β*, allowing node ranks to vary while keeping the same tree shape. For small values of *α* (in particular *α* < 0), the smallest subintervals have a higher probability of being selected, so smaller clades tend to be older. On the contrary, for large values of *α*, the largest subintervals have a higher probability of being selected, so smaller clades tend to be younger. We notice that as *β* gets close to –2 the effect of *α* vanishes, since at all times there is merely one edge that can split. In maximally unbalanced tree shape (*β* = –2), there is only one ranked tree shape and the order of nodes is fixed, so *α* plays no role.

As is well-known, the tree obtained with *β* = 0 has the same shape has the tree generated with the Yule process (Yule 1925) or the Kingman coalescent (Kingman 1982) after ignoring node ranks (Nee 2006; Lambert and Stadler 2013). When *α* = 1 in addition to *β* = 0, we show in Appendix 1 (Proposition 1), available in Supplementary Materials, that our model generates the same tree shape with node ranks as Yule trees, which is actually known to be the same as the ranked tree shape of the Kingman coalescent tree.

The version of the model we present here only allows simulation of trees with *β* > –1, as the Beta distribution is only defined for positive parameter values. Actually, our model coincides with the ranked tree in a self-similar, binary fragmentation with self-similarity index *α* and with fragmentation measure 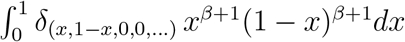 (as defined in Bertoin 2002, 2006), which makes sense as soon as *β* > –2. In Appendix 1 (see in particular Proposition 3), we present an algorithm based on fragmentation processes equivalent to that presented above (using one additional approximation parameter e, consistently set to 0.001). Albeit less intuitive, this method allows us to simulate trees for all *β* > –2.

Last, it is important to notice that our model is both exchangeable and sampling consistent. It is exchangeable because labels can be swapped without changing the distribution of the tree, since marks all have the same distribution. It is sampling consistent because removing tip labelled *n* + 1 (or any other tip, by exchangeability) amounts to removing mark *U*_*n*+1_, which does not modify the ranked tree shape obtained from marks (*U_i_*)_*i*∈{1,…,*n*}_.

### Incorporating non-random extinctions

In order to map each clade of our random phylogeny to its frequency (*i.e*., relative abundance or relative range size), we add, into a second version of the model, a new parameter *η* ≥ 0. Each time an interval *X* is split into two subintervals, *X_left_* and *X_right_* with widths |*X_1eft_*| = R |*X*| and |*X_right_*| = (1 – *R*) |*X*|, each of the two subtrees is granted a part of the abundance *A_X_* of the parental clade equal to

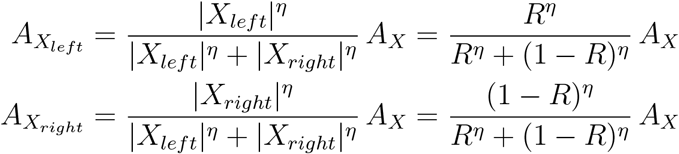

This way of allocating frequencies to taxa is reminiscent of the ‘broken stick model’ (MacArthur 1957; MacArthur and Wilson 1967; Colwell and Lees 2000), where the unit interval is broken into subintervals each representing the frequency or resource share of each species or clade in the community. This is usually done by throwing uniform points independently in the interval or by throwing the points sequentially, always to the right of the last one, leading to the Poisson-Dirichlet distribution appearing in mathematical population genetics (Feng 2010; Ewens 2012) as well as in the neutral theory of biodiversity (Hubbell 2001).

The model remains sampling-consistent insofar as each *A_X_* is interpreted as the abundance of a whole clade, that is the sum of abundances of all species belonging to this clade, present or not in the sample. Sampling consistency now means that generating a ranked tree shape with relative abundances on *n* tips is equivalent to the following process: generate a ranked tree shape with relative abundances on *n* +1 tips, remove one tip at random and sum the abundance of the removed tip to that of its sister clade (i.e., the clade descending from the interior node connected to the removed tip by a pendant edge).

If *η* = 1, then *A_X_* = |*X*| so that each clade is granted an abundance that is in mean proportional to its richness, which means each tip gets the same abundance on average. When *η* > 1, the largest of the two daughter clades gets a share of the abundance that is (in mean) more than its share in species richness, so species in large clades tend to be more abundant than species in small clades; the opposite happens for *η* < 1, and species in small clades tend to be more abundant than species in large clades. Variance in species abundances increases with |*η*|. Simulations of species relative abundance (or range) distributions are shown for different values of *η* in Figure 3 and in Appendix 1.

**Figure 3:**
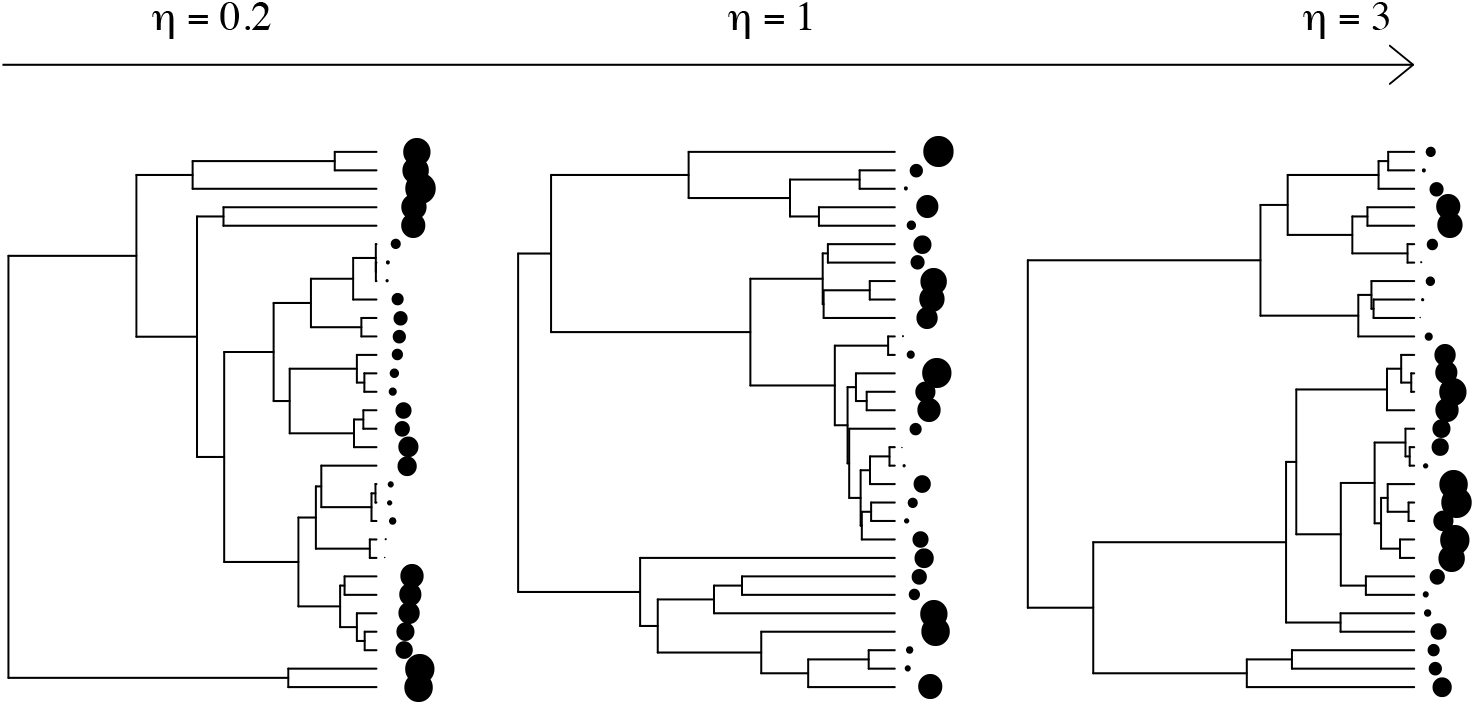
Distribution of species frequencies across the tips of phylogenetic trees for different values of *η* (correlation clade size-frequency). Dot sizes sort species according to their frequency (larger dots for more abundant species). Parameter values: *η* = 0.2, 1 or 3 (from left to right), *β* = 0, *α* = 0, number of species *N* = 30, *ϵ* = 0.001. Results with *β* = –1.9 are shown in the online Appendix 1 available as Supplementary Material.

In the extinction numerical experiment, we determine the order of species extinctions deterministically based on their rank in abundance: the rarer species are the first ones to go extinct, whereas more frequent species go extinct last (Fig. 3). The case *η* = 1 where each tip gets the same abundance on average is roughly equivalent to the field of bullets model of extinction (see Proposition 2 in Appendix 1; in the case *β* = –1, the equivalence is exact). This modeling approach allows us to tune the sign and strength of the correlation between the richness of a clade and the extinction risk of its species.

### Testing the effect of β, α and η on PD loss

The effect of all three model parameters on the relationship between species loss and PD loss is studied in a systematic way by simulation. We considered values of *β* in (–2,10], values of *α* in [–3, 3] and *η* in [0.1, 3]. Because our model specifies how interior nodes are ranked in time but not their actual timing, we use a pure-birth process to generate node depths, adding the latter on top of ranked tree shapes. The use of another model for generating node depths leads to qualitatively similar results, albeit quantitatively different (as an illustration, we show results with edge lengths set as in the Kingman coalescent in Appendices 4 and 6, available as Supplementary Material).

For each set of parameter values, we generated one hundred trees with one hundred tips (*N* = 100). We sequentially removed extinct species from these trees (in the order of increasing species abundances, as explained earlier), and computed the remaining PD (sum of all branch lengths; Faith 1992) for increasing fractions of extinct species.

### Parameter inference

We infered the parameters *β, α* and/or *η* from simulated or empirical datasets by maximum likelihood. As is already well-known (Aldous 1996; Blum and François 2006; Lambert et al. 2017), the likelihood of a labelled tree shape under Aldous’ *β*-splitting model is explicit. Since the likelihood of the tree shape under our model is the same as in Aldous’ model (and in particular independent of *α* and *η*) we can use it to estimate *β*. In contrast, computing the likelihood of the ranked tree shape requires to follow through time the lengths of all intervals of the partition containing marks, which may decrease without separating marks (unsampled species). Given that the likelihood of the ranked tree (with or without tip abundances) with the additional knowledge of interval lengths is explicit, we use a Monte-Carlo data augmentation procedure, in which the augmentation data are the numbers and sizes of unsampled splits on each branch (which allow us to reconstruct the interval lengths through time). The likelihood of the ranked tree with tip abundances is then computed by averaging over augmentations, and is optimized over possible values of (*α, η*).

We first tested our ability to infer the model parameters on simulated trees. To do so we simulated trees with 20, 50 and 100 tips for all possible combinations of *α* in {–1, 0,1, 2}, *β* in {–1, 0,1} and *η* in {0.2, 0.5,1,1.5, 2}. For each tree size and parameter combination, we simulated 20 trees with tip abundances, for a total number of 3600 trees.

We then inferred the model parameters on these trees and compared them to the values used in the simulations. The inference of the parameter *β* was straightforward, being computed as the maximum likelihood estimate on the interval] –2, 10] with the `function maxlik.betasplit` from the R-package `aapTreeshape` (Bortolussi et al. 2006). The parameters *α* and *η* were estimated with the method introduced hereabove, with values respectively constrained on the interval [–4, 4] and [0.1,10]. The value of *η* (minimum size of unsampled splits, see Appendix 1 in the Supplementary Materials) was here again fixed to 0.001.

The validation of this estimation procedure allowed applying it to real bird family trees. We used the MCC tree from Jetz et al. (2012), and pruned it to keep family level phylogenies. We kept only the phylogenies that included at least 50 species, and used range sizes from Map of Life (https://mol.org/) as tip data. The value of ∊ and the constraints on parameter ranges were here the same as in the test on simulated phylogenies.

The model was coded-and the analyses of phylogenetic trees were performed-using `R` (R Development Core Team 2012) and the R packages `cubature` (Johnson and Narasimhan 2013), `ape` (Paradis et al. 2004), `sads` (Prado et al. 2015), `apTreeshape` (Bortolussi et al. 2006) and `picante` (Kembel et al. 2014).

## Results

### Influence of ranked tree shape on PD loss

Here we only address the influence of *α* on PD loss, assuming a field of bullets model for species extinctions (*η* = 1). The expected PD loss is then a convex function of the fraction *p* of extinct species (as proved mathematically for any binary tree under the field of bullets model, see Eq (34) in Lambert and Steel 2013), always lying below *p* (Fig. 4.A,C,E,G).

**Figure 4:**
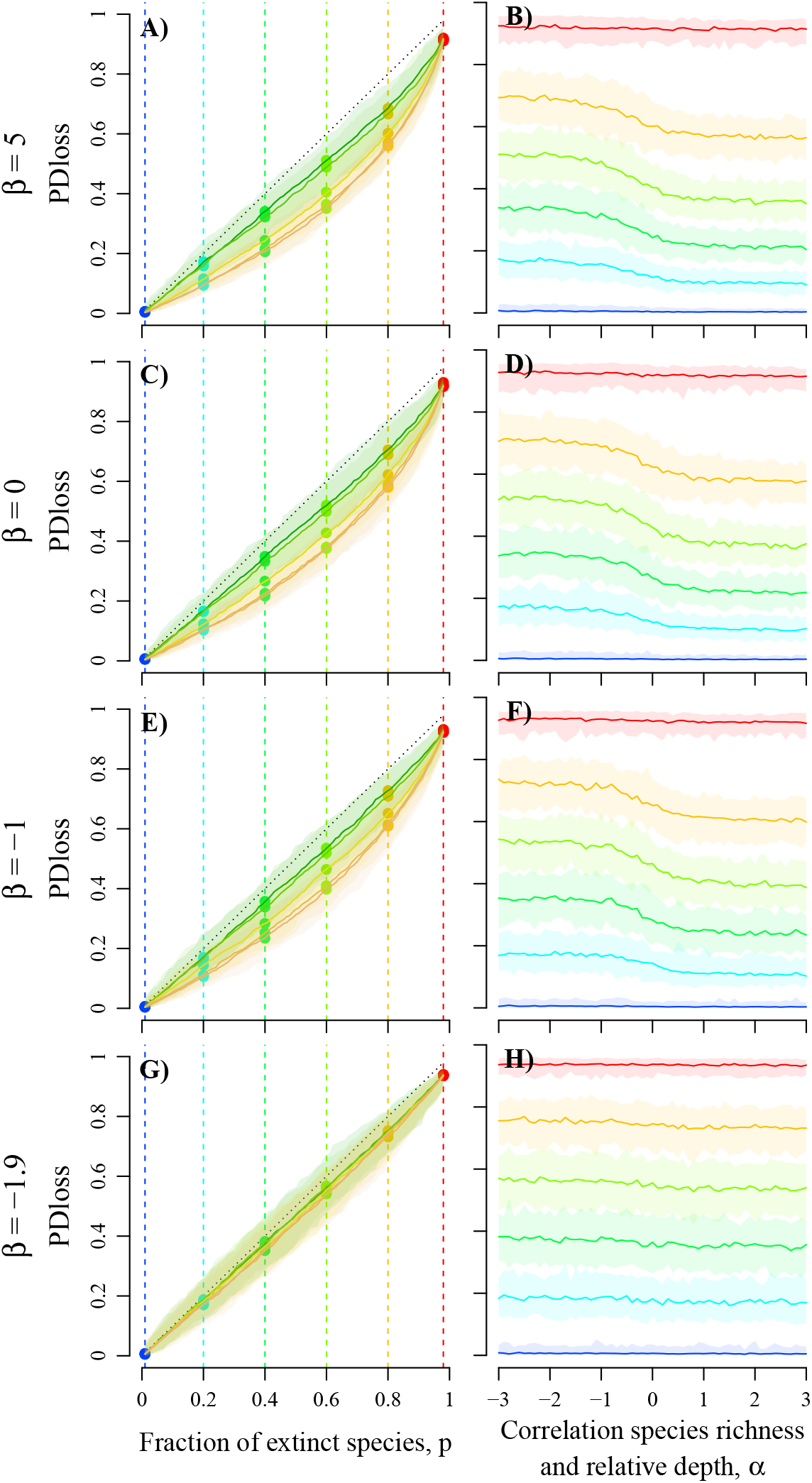
Influence of the ranked tree shape (tree balance *β* and correlation clade size-age *α*) on philogenetic diversity (PD) loss, for increasing fractions of species extinctions. Tree balance *β* changes from 10 (top row, ‘bush trees’) to –1.9 (bottom row, ‘comb trees’). Results are shown either as a function of the extinction fraction p (left column; for different *α* values) or as a function of *α* (right column; for different extinction fractions *p*). Extinction fraction *p* increases from 0.01 to 0.98 (from left to right in A, C, E, G; from blue to red in B, D, F, H). The dotted lines in A, C, E, G show the bisector. Results are based on 100 simulation replicates: plain lines give median values and light areas give 95% confidence intervals. Other parameter values: number of species *N* = 100, *ϵ* = 0.001.

Consistently with previous studies (Nee and May 1997; von Euler 2001) we find that when the relation between depths and richnesses of clades is similar as that in Yule trees (*α* = 1), very unbalanced trees (comb-like trees) lose more PD in the face of species extinctions than Yule or more balanced trees (Fig. 4. G-H *vs*. A-D, with *α* = 1). The effect is non-linear in *β*: the tree shape has little influence on the loss of PD when *β* ≥ –1, but increases sharply as *β* decreases from –1 to –1.9 (results as a function of *β* in Appendix 2, available as Supplementary Material). Unbalanced tree shapes are associated with the presence of long edges leading to evolutionary distinct species (Fig. 2). These edges constitute an important fraction of the phylogenetic diversity in unbalanced species trees, so that their extinction generates a significant drop in PD. As *β* gets closer to –2 (case of the ‘comb tree’), the expected PD loss approaches the fraction of extinct species (Fig. 4.G).

Considering ranked tree shapes shows, however, that the order of nodes has a significant influence on the loss of PD, and on the effect of *β* on this loss. If the depth and richness of clades are positively correlated (*α* > 0), the loss of PD is reduced, especially at intermediate extinction fractions (Fig. 4.A-F). This is because the smallest subtrees, more prone to early extinction, are younger and hence contain a lower fraction of the phylogenetic diversity (Fig. 2). If the depth and richness of clades are negatively correlated (*α* < 0), the loss of PD rises, especially at intermediate extinction fractions. The smallest subtrees, prone to extinction, are older and hence contain more evolutionary distinct species (Fig. 2). This generates losses of PD similar to those observed when the tree shapes are very unbalanced (PD loss equal to the fraction of extinct species).

As expected, the effect of *α* is evened out in very unbalanced trees (*β* close to –2; Fig. 4.G-H), for which the loss of PD remains close to its highest value whatever the value of *α*. In the case of the maximally unbalanced tree shape, there is only one ranked tree shape and the order of nodes is fixed.

All these effects of ranked tree shapes on the loss of PD are qualitatively conserved if node depths are distributed as in the Kingman coalescent (instead of the Yule process). In the case of Yule trees, PD loss slightly increases with the initial size of the tree, an effect which is due to more efficient sampling of large values in the common (exponential) distribution of node depths. Yet the results presented above are qualitatively conserved if the size of phylogenetic trees changes (analyses performed with number of species *N* = 50 and *N* = 200; see the online Appendices 3 and 4 available as Supplementary Materials).

### Influence of non-random extinction risks on PD loss

Correlations between the richness of a clade and its relative abundance (here directly influencing the extinction risk of its species) may have a paramount influence on the loss of PD in the face of extinctions (Fig. 6). In trees with ranked tree shapes similar to Yule trees (*β* = 0, *α* = 1), the concentration of high extinction risks in small clades (*η* > 1) increases the loss of PD, by promoting the extinction of entire clades (Fig. 5). In contrast, when extinction risks are higher in larger clades (*η* < 1), phylogenetic redundancy (and hence the likelihood of conserving at least one species per subtree) limits the loss of PD until high extinction levels.

**Figure 5:**
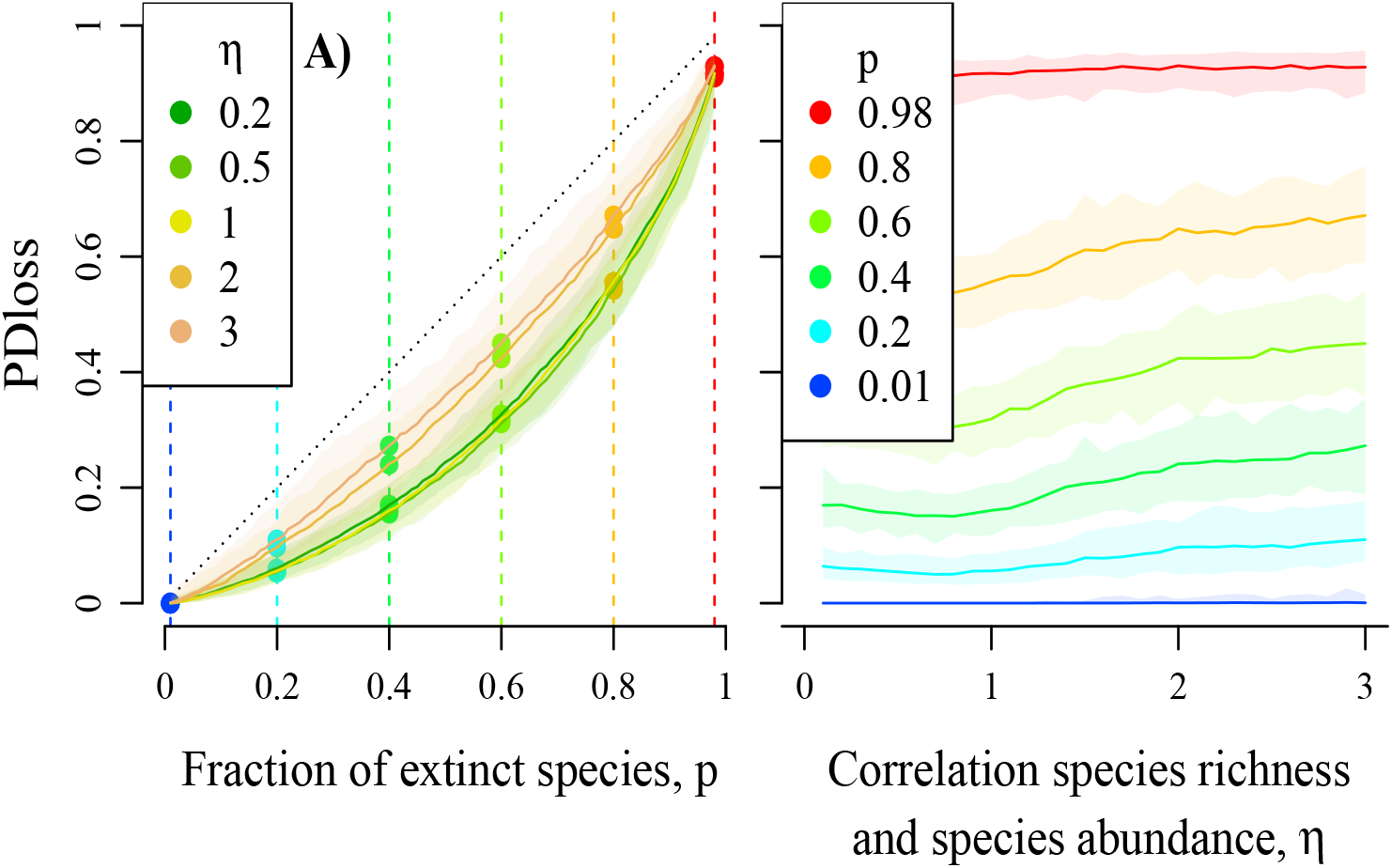
Effect of *η* (correlation clade size-frequency) on PD loss in Yule trees, for increasing fractions of species extinctions *p*. Results are shown either (A) as a function of the extinction fraction *p* (for different *η* values, with dotted lines showing the bisector) or (B) as a function of *η* (for extinction fractions *p* increasing from 0.01 to 0.98 from blue to red). Results are based on 100 simulation replicates: plain lines give median values and light areas give 95% confidence intervals. Parameter values: *β* = 0, *α* = 0, number of species *N* = 100, *ϵ* = 0.001.

**Figure 6:**
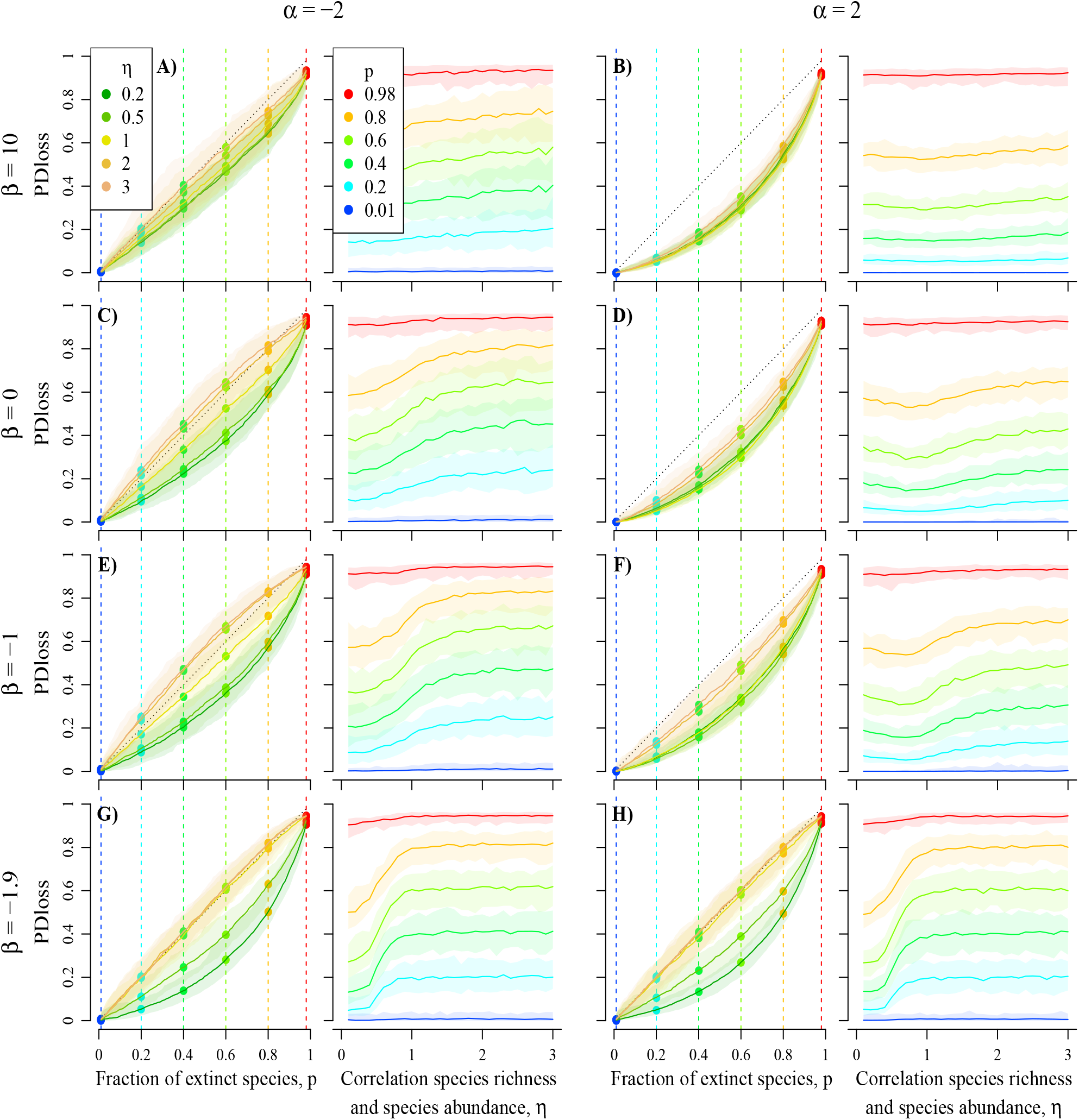
Effect of *η* (correlation clade size-frequency) on PD loss, for different ranked tree shapes and increasing fractions of species extinctions. Tree balance *β* ranges from 10 (top row, ‘bush trees’) to –1.9 (bottom row, ‘comb trees’), and correlation clade size-age *α* ranges from –2 (A, C, E, G) to 2 (B, D, F, H). Results are shown either as a function of the extinction fraction p (left side; for different *η* values, and with dotted lines showing the bisector) or as a function of *η* (right side; for extinction fractions *p* increasing from 0.01 to 0.98 from blue to red). Results are based on 100 simulation replicates: plain lines give median values and light areas give 95% confidence intervals. Other parameter values: number of species *N* = 100, *ϵ* = 0.001.

The effect of *η* is modified by the ranked shape of species trees. Correlations between clade richness and clade depth (set by *α*) modulate the additional loss of PD induced by *η* > 1 (i.e. lower abundances in smaller clades; Fig. 6.A-F). When *α* < 0,smaller clades are not only more prone to extinction but also have deeper nodes, hence more evolutionary distinct species, which increases even further the loss of PD. Unlike in the field of bullets model, the expected PD loss as a function of the fraction p of extinct species can even change from convex to concave, and so take values larger than p (Fig. 6C,E). When *α* > 0, smaller clades are more prone to extinction but have shallower nodes, which counteracts the increase of PD loss due to *η* > 1. To summarize, PD loss is increased when *η* > 1 compared to *η* = 1, with a maximal effect for negative values of *α*, progressively flattening as *α* grows.

We call ‘thin ice zone’ the region of parameters corresponding to the theoretical phylogenies that suffer a maximal rate of PD loss straight from the first few extinction events, that is, close to 1% of PD lost for the first 1% of species lost. In the plane (*α, η*), the ‘thin ice zone’ correspnds to {*α* < 0, *η* > 1}. As testified by Fig. 6, phylogenies in this zone can even suffer a rate of PD loss which is larger than 1 from the first extinction and sustains itself above 1 throughout the extinction crisis.

In contrast, *α* has little effect on the decrease in PD loss induced by *η* < 1 (*i.e*., higher abundances in small clades). Indeed, when *η* < 1, the deepest nodes are always protected regardless of the value of *α*: when *α* < 0 the deepest nodes are in small clades which are protected from extinctions by their high relative abundances (due to *η* < 1); when *α* > 0, the deepest nodes are in large clades which are protected by phylogenetic redundancy.

The influence of *η* on PD loss is amplified by unbalanced tree shapes (*β* < 0; Fig. 6.E-H) and buffered by balanced tree shapes (*β* > 0; 6A-B), because lower values of *β* enhance richness inequalities between clades and raise in turn the influence of *η* on PD loss. This interaction between parameters *η* and *β* overwhelms the influence of *α* (Fig. 6). In the plane (*β, η*), the ‘thin ice zone’ is {*β* < –1, *η* > 1} and the previous remark thus implies that in the three-dimensional parameter space, the thin ice zone is {*α* < 0 or *β* < –1; *η* > 1}.

Interestingly, the effect of *β* is highly dependent on how extinction risks are distributed within the phylogeny (Fig. 7, and results with other *α* values in Appendix 7 available as Supplementary Material). For *η* = 1, we recover the well-known pattern of decreased PD loss as the tree gets more balanced. However, for *η* < 1 we see the reverse pattern, that is PD loss increases with the balance of the tree. Recall that *η* < 1 buffers PD loss, because extinction risks are clustered in the bigger clades which also display higher phylogenetic redundancy (smaller pendant edges). When the tree is maximally unbalanced, *η* < 1 causes the longest pendant edge to subtend the tip with the largest abundance (and hence to be the last to become extinct). Therefore, the order of extinctions coincides exactly with the increasing order of pendant edge lengths, which results in minimal PD loss for any given level of extinction. In a more balanced phylogeny, the distribution of clade sizes is more even and the buffering effect of the clustered extinction on PD loss is reduced.

**Figure 7:**
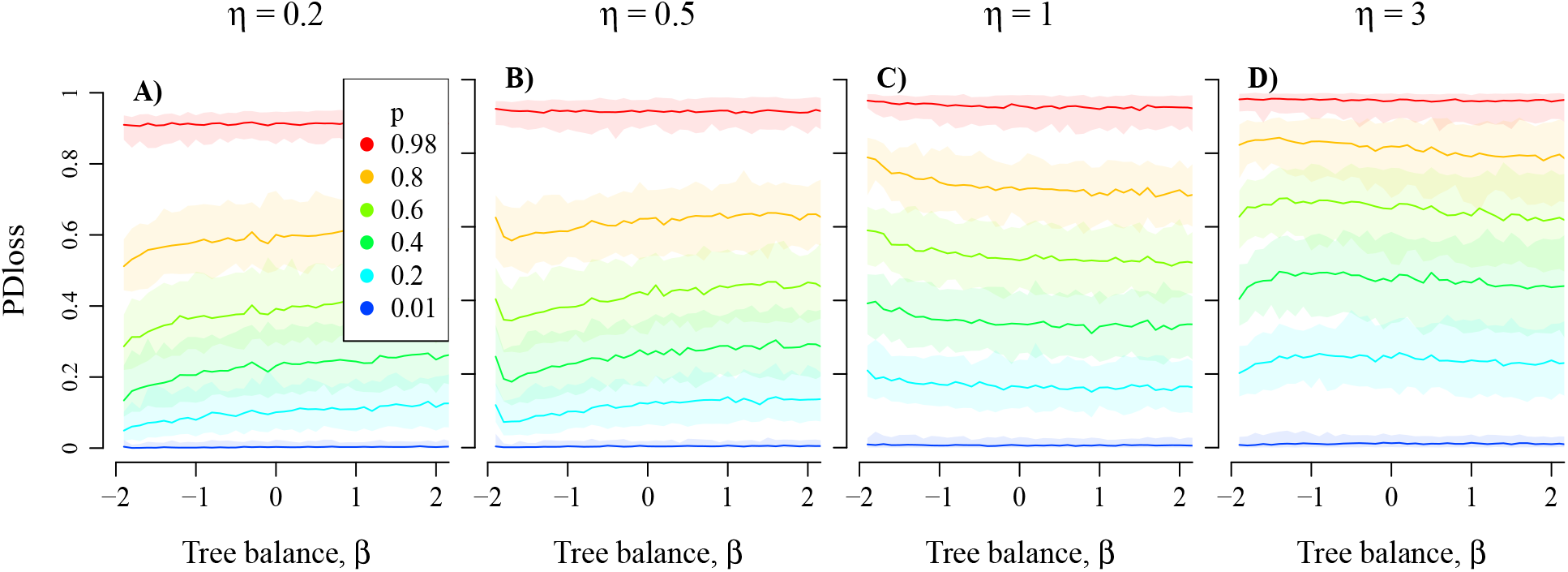
Effect of tree balance *β* on PD loss, for different correlations clade size-frequency *η*, and increasing fractions of species extinctions *p*. The correlation clade size-frequency *η* ranges from 0.2 (left) to 3 (right), and the extinction fraction p increases from 0.01 to 0.98 (from blue to red). Results are based on 100 simulation replicates: plain lines give median values and light areas give 95% confidence intervals. Other parameter values: clade size-age *α* = –2, number of species *N* = 100, *ϵ* = 0.001.

For *η* > 1 we again recover the well-known pattern of decreased PD loss with increasing *β*. However, when we also have *α* < 0, the relationship between PD loss and *β* is not monotonic, that is for any particular level of extinction, the maximal PD loss is reached for trees with intermediate balance. Recall that *α* < 0 causes small clades to be relatively older and so to contribute more to PD. The maximal loss of PD thus occurs when extinction risks cluster in small clades. And indeed, when *η* > 1, at each splitting event the species-richer subtree gets a bigger abundance than the species-poorer subtree. However, within a given clade, the abundance of a species should decrease with the number of nodes (splitting events) on its lineage. This latter effect is stronger in unbalanced trees; in balanced trees, extinction risks cannot cluster in small clades, due to the absence of small clades. Trees with intermediate balance do display small clades, and these small clades are large enough to share their low abundance (*η* > 1) into a few species with very low abundance. These species go extinct first, resulting in maximal PD loss.

### Effect of species extinctions on tree shape

We study the effect of species extinctions on tree shape, seeking in particular to check if the influence of *η* on the patterns of PD loss can be explained by changes in tree shapeas species go extinct. Fig. 8 shows the imbalance (defined here as the maximum likelihood estimate *β* of the parameter *β*) of the species tree computed after a fraction *p* of its species have become extinct. When *η* = 1, tree balance is very little altered by extinctions except in very balanced trees, as predicted by the sampling consistency of the model (*η* = 1 amounts to removing species at random except when *β* ≫ 1, see Appendix 1). When *η* < 1, trees tend to become more and more balanced as *p* increases (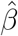 increases with *p*), whereas when *η* > 1 trees tend to become more and more similar to Yule trees (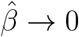 as *p* → 1). The effect of *η* on PD loss cannot be reduced to its effect on changes in tree shape due to extinctions. On the one hand, *η* mostly affects the shape of trees with *β* > –1 (Fig. 8), whereas tree shape has most effect on PD loss when *β* varies between –2 and –1 (Fig. 4.A,C,E with *α* = 0). In addition, if the effect of *η* on tree shape had a significant influence on PD loss, *η* > 1 should increase this loss when *β* > 0 (by decreasing the balance of trees; Fig. 8.D) and decrease it when *β* < 0 (by increasing the balance of trees). Yet, the changes we observe in the effect of *η* > 1 on PD loss for different *β* values are the reverse of this prediction. Therefore, the indirect effects of *η* (through changes in tree shape) are negligible compared to its direct effects (through non-random distribution of extinction risks).

**Figure 8:**
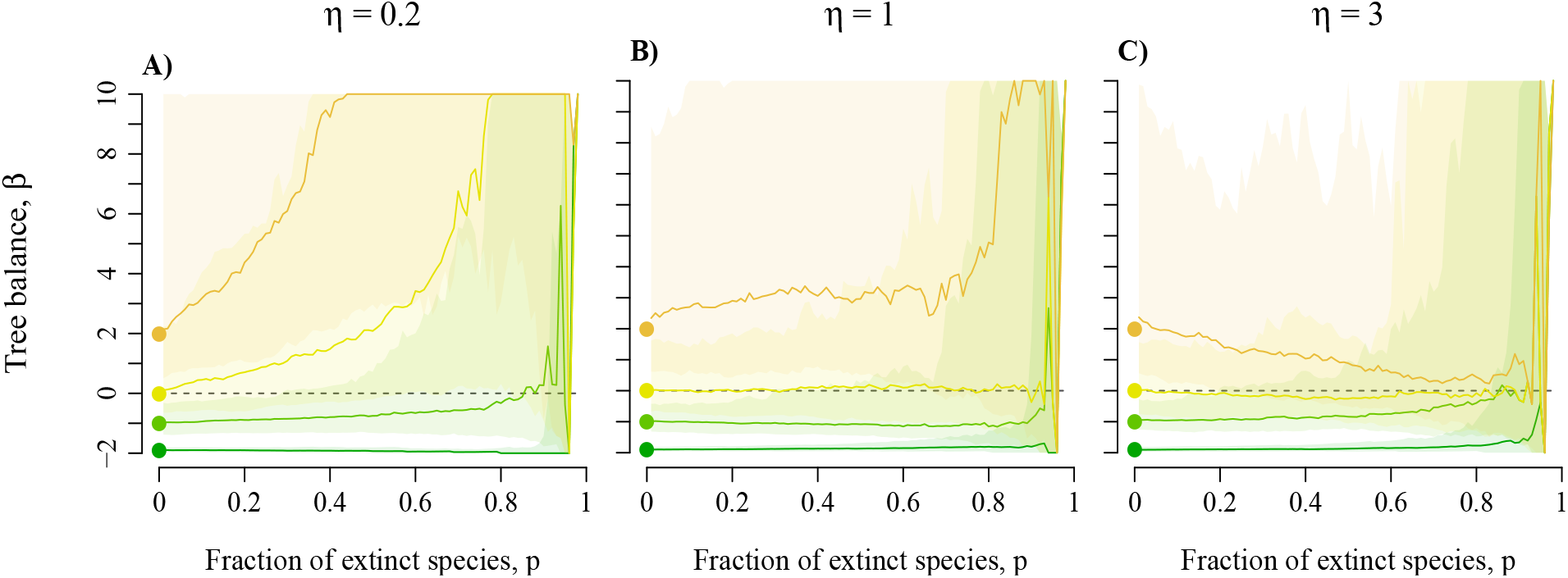
Effect of *η* (correlation clade size-frequency) on the balance of phylogenetic trees after extinctions (MLE 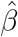 of *β*). Initial tree balance *β* ranges from 10 (brown dots and lines, ‘bush trees’) to –1.9 (green dots and lines, ‘comb trees’). Extinction fraction p increases from 0.01 to 0.98 (from left to right). Results are based on 100 simulation replicates: plain lines give median values and light areas give 95% confidence intervals. Other parameter values: number of species *N* = 100, *ϵ* = 0.001, *α* = 0.

As precedently results on the effects of non-random extinctions on the loss of phylogenetic diversity are conserved when node depths are distributed as in the Kingman coalescent, or when the size of phylogenetic trees changes (analyses performed with *N* = 50 and *N* = 200; see the online Appendices 5 and 6, available as Supplementary Material).

### Parameter inference

When tested against simulated data, the Monte-Carlo inference algorithm by data augmentation performs reasonably well on phylogenies with more than 50 tips for a wide range of parameters (see the online Appendix 8, available as Supplementary Material). As expected, the estimation of *β* on trees with at least 50 tips is accurate, since the likelihood formula of the unranked tree is explicit, and this accuracy increases as *β* decreases. The inference algorithm also returns overall good estimates of *η* and *α* whenever *η* > 0.3.

The inference of *α* is unbiased except in the cases where *β* < 0 and *η* < 0.3. This corresponds to cases where he unsampled nodes are numerous because *β* is small, and they have a strong impact on the reconstruction of intervals because *η* is small. The inferred *η* is overestimated for trees with only 50 tips. For *β* < 0 and *α* ≥ 0, *η* is sligthly overestimated whatever the tip number. For *β* > 0 and *α* ≤ 0 inferences are good for trees with at least 100 tips.

### Empirical values

Estimates of parameter values on real data shows consistent patterns across all bird family trees. Unsurprisingly, we find negative *β* values, mostly comprised between 0 and -1, hence corresponding to unbalanced trees (see online Appendix 9, available as Supplementary Material). Since the estimation of *β* is quite accurate for low true values of *β* and is biased towards larger estimates than the true value otherwise, these estimates can be taken with confidence. The estimates of *η* vary between 1 and 1.5. This indicates that, within bird families, species in small clades tend to have smaller range sizes than species in larger clades. The above study showed that low *η* values can be difficult to detect in unbalanced trees. Yet when this is the case, *η* is found to be close to the maximal value allowed in the inference (here 10), which is not the case here. We can therefore be confident that these values do not reflect a biais in the inference, but reflect a true pattern in the distribution of range sizes within the phylogenies. Finally, the estimates of *α* are clustered around 0, indicating that there is no correlation between clade size and clade depth within each bird phylogeny. This in contrast with what is expected in most explicit models of diversification, where larger clades take more time to diversify, resulting in a strong positive correlation between the depth and the size of clades.

When jointly infering of *α* and *η* the choice to use range size to infer *η* is likely to have an impact on the inferred *α* (because the values of the intervals are reconstructed using tip values, unappropriate tip values would lead to uncorrect *α*). Therefore we also ran the inference of *α*: we wind fairly similar results between values obtained with the inference of alpha only compared to the full inference (the median of the inferred *α* for trees with at least 50 tips is 0.19 when *α* is inferred alone and 0.05 when both *α* and *η* are inferred), indicating that tree shape is indeed driving the result (online Appendix 9 available as Supplementary Material equivalent as Fig. 9 with the *α* inferred without knowledge of the tip range sizes).

**Figure 9:**
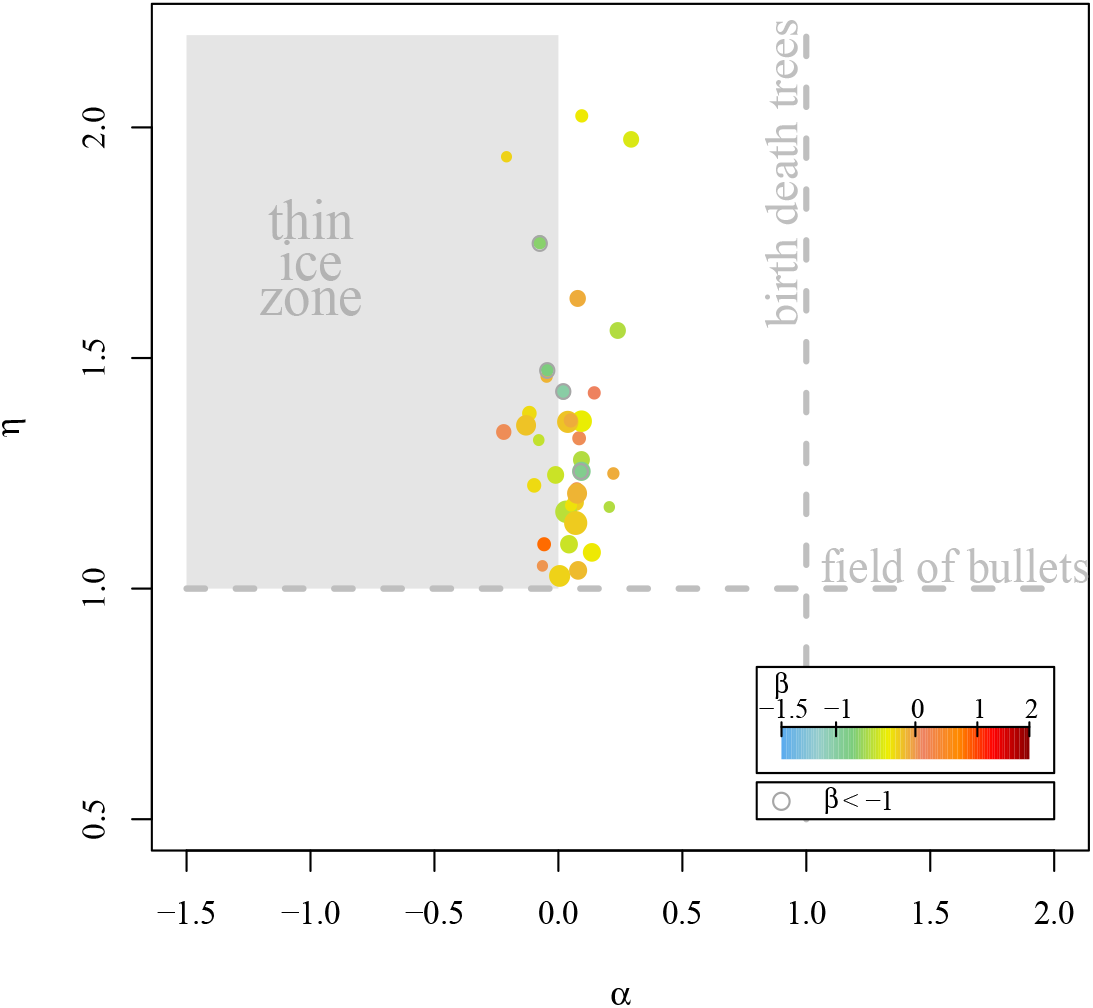
Inferred model parameters on bird family trees of 50 tips or more. *α* maximum posterior estimate (x-axis), *η* maximum posterior estimate (y-axis) and *β* maximum likelihood estimate (point color). Point sizes are proportional to the number of tips in trees, *N*. The dashed vertical line shows the value of *α* for trees generated by a birth-death model, and the dashed horizontal line shows the value of *η* for which extinction probabilities are distributed within the tree as in a field of bullets model. For all inferences, *ϵ* was set to 0.001.

## Discussion

### A new integrative measure of correlation between clade depths and sizes, α

We introduced here a new model for random ranked tree shapes with a fixed, arbitrary number of tips. This model features two parameters, *β* and *α* tuning respectively the shape of the tree and the order of its nodes. Trees with *β* ≤ 0 are imbalanced and trees with *β* > 0 are balanced. Whatever the value of *α*, the shape of the tree is the same as in Aldous’ *β*-splitting model (Aldous 1996, 2001). Large clades coalesce deep in the tree when *α* > 0 and are shallower than smaller clades when *α* < 0. When *β* = 0 and *α* = 1, the tree has the same ranked shape as the Kingman coalescent and the Yule tree. In addition, this model is the first model (except the two aforementioned models and the trivial case of the ‘comb tree’) for ranked tree shapes satisfying sampling-consistency, in the sense that a tree with n tips has the same distribution as a tree with *n* + 1 tips with one tip removed at random. This property is essential to ensure the robustness of the model with respect to incomplete taxon sampling (Heath et al. 2008; Cusimano et al. 2012; Stadler 2013).

Predictions from this model highlight the importance of accounting for node ranks to understand forthcoming changes in macroevolutionary patterns of phylogenetic diversity. They show in particular that the relationship between the species richness of a clade and its relative depth in the tree, set by parameter *α* in the model, can have profound impacts on the rate of PD loss (Fig. 4). This parameter *α* constitutes a new index quantifying the relationship between the depth and size of clades. A large number of studies already looked at the depth-size correlation, assessing its existence (significance, sign and pattern) across multiple phylogenetic trees-based on one value of species richness and crown or stem age per phylogeny (*e.g*., Magallon and Sanderson 2001; Bokma 2003; Ricklefs 2006; McPeek and Brown 2007; Rabosky et al. 2007; Ricklefs 2007a; Rabosky 2009; Rabosky et al. 2012). These studies notably aimed at testing the hypothesis of time-limited diversity patterns, versus hypotheses of diversity set by diversification rates or by limits to diversity (McPeek and Brown 2007; Ricklefs 2007b, 2009; Rabosky 2009; Barraclough 2010; Rabosky 2013). Our new index *α* is different in that it can be measured by maximizing the likelihood on a single phylogeny, implicitly integrating over all subclades of this phylogeny. An interesting consequence is that one does not have to choose which clades to include in the analysis. For example, *α* is not sensitive to the definition of higher taxa (Stadler et al. 2014). Moreover, similarly to the index *β* (compared to other measures of tree imbalance; Kirkpatrick and Slatkin 1993; Aldous 1996, 2001), *α* is a measure of depth-size correlation computed as the maximum likelihood estimate of a model-based parameter. Last, we stress that our model does not require the precise knowledge of node datings in the phylogeny but only the relative positions of nodes in time, which preserves *α* alpha estimates from the inaccuracies of time calibrations (Kumar 2005; Welch and Bromham 2005; Pulquério and Nichols 2007; Forest 2009; Schwartz and Mueller 2010).

### Ranked tree shapes and the loss of phylogenetic diversity

Our results confirm that in the field of bullets model unbalanced trees undergo stronger loss of PD than balanced trees, under equal fraction of species extinctions. This property was already well-known (Nee and May 1997), but is important to recall given the predominance of unbalanced phylogenetic trees in nature (*β* values being often close to –1; *e.g*., Guyer and Slowinski 1991; Heard 1992; Guyer and Slowinski 1993; Slowinski and Guyer 1993; Mooers 1995; Purvis 1996; Mooers and Heard 1997; Blum and François 2006). However, our results also show that the temporal order of nodes among subtrees (set by the parameter *α*) may have even stronger effects than their distribution among subtrees (set by parameter *β*; compare the effect of the latter in Appendix 2 to that of *α* on Fig. 4). Besides, *α* values below 0 cause drops of PD almost as abrupt as those observed with ‘comb’ shapes (*β* close to –2, with *α* = 1; Fig. 4.D,H). It is therefore essential to consider the ranked shapes of species trees to understand the expected patterns of loss of phylogenetic diversity.

Values of *α* deviating from 1 may arise from differences in stages of diversification among subtrees, resulting from heterogeneity in biotic or abiotic factors acting on diversification processes in different parts of the species tree. This could be due to bursts of diversification in certain subtrees (*e.g*., following from key innovations or from migration to empty spatial or ecological space), either recently (resulting in *α* < 0) or early in the history of clades (resulting in *α* > 0). Alternatively, *α* values deviating from 1 could be linked to changes in extinction rates in distinct parts of the tree (*e.g*., due to changes in the biotic or abiotic environment of phylogenetically related species sharing similar ecological niches). Age-dependent speciation (Hagen et al. 2015) and extinction (Alexander et al. 2015) are also likely to make node ranking deviate from what is expected in a homogeneous birth-death model. Heterogeneity in diversification rates across the species tree associated with asymmetric competition among species (*e.g*., evolutionary advantage to previously established species) could limit diversification in younger subtrees, hence leading to *α* > 0. Last, *α* can be found negative due to the presence of relictual lineages, *i.e*., old clades harboring few species surviving to the present.

### Modeling non-random extinctions: η and the loss of phylogenetic diversity

The incorporation of parameter *η* within the framework provided by Aldous’ *β*-splitting model allowed us to go beyond the field of bullets assumption. In passing, we devised a model of abundance distributions (equivalently interpreted as range size distributions) covarying with the phylogeny, in the broken-stick tradition (MacArthur 1957; MacArthur and Wilson 1967). When *η* > 1, the most abundant species are in species-rich clades whereas when *η* < 1 the most abundant species are in species-poor clades. When *η* = 1 all species have the same abundance on average. Here, extinctions are assumed to occur sequentially in the order of increasing abundances. In nature, relative extinction risk indeed depends on species frequency, but also on many other features (*e.g*., dynamics of population growth or decline, fragmentation into subpopulations, biotic or abiotic changes; IUCN 2012), and may have a significant stochastic component. The simple framework we use to determine extinctions allows us to focus on the direct impact of the distribution of ranked abundances within trees on the loss of phylogenetic diversity. This framework can easily be modified to include extrinsic causes of extinctions.

Previous studies concluded that PD loss is increased if extinction risks are clustered in the phylogeny (Davies and Yessoufou 2013), but that this effect is not substantial (Parhar and Mooers 2011). Our model shows that the effect on PD loss depends on the way these extinction risks are distributed among clades: PD loss is increased by *η* > 1 (*i.e*., higher extinction risks in small clades; Fig. 6). Such a distribution of extinction risks may arise from subtrees having low species richness because of higher extinction rates, either due to intrinsic factors (species features that would make them more susceptible to extinction; *e.g*., long generation time, or low variance or phenotypic plasticity of key ecological traits providing resistance to perturbations or evolutionary advantages in relation to biotic interactions; Purvis et al. 2000c; Johnson et al. 2002) or to extrinsic factors (threats affecting the spatial or ecological space shared by species of the subtree; *e.g*., Russell et al. 1998; Hughes 1999; Purvis et al. 2000c; von Euler 2001; Johnson et al. 2002). Higher extinction risks in small subtrees could also be due to resource limitation affecting simultaneously the density of individuals and the diversity of species, and hence demographic stochasticity; or to stabilizing selection (*e.g*., due to competition or to the absence of available spatial or ecological space in the surrounding environment), limiting adaptation and increasing species vulnerability in the face of perturbations (Purvis et al. 2000c; Purvis 2008).

In contrast, *η* < 1 buffers the loss of phylogenetic diversity. Higher extinction risks in larger subtrees could result from a trade-off between species richness and average species abundance, provided constrained metacommunity size (with variation along this trade-off following for instance from landscape structure and dynamics, such as geographical isolation affecting the occurrence of allopatric speciation events), from recent speciation events associated with a decrease in average species abundance, geographical range or niche width, or from recent extinction events that removed the most extinction-prone species from certain clades (leaving the latter smaller and with less extinction-prone species; Schwartz and Simberloff 2001; Lozano and Schwartz 2005).

Hence, *η* is expected to vary across clades according to the metacommunity structure and the underpinning diversification dynamics. Given its striking effects on PD loss, this factor should also be accounted for to understand potential future losses of phylogenetic diversity.

### Combined effects of β, α and η: reversing some expected patterns of PD loss

The influence of *η* on the loss of PD is enhanced by *α* < 0 (small clades containing evolutionary distinct species) and *β* < 0 (more variability in clade richness) (Fig. 6). However, a stronger clustering of extinction risks does not necessarily lead to higher loss of PD (*e.g*., if extinctions occur first in richer subtrees–which contain more phylogenetic redundancy–as in the case when *α* > 0 and *η* < 1).

These interactions between the effects of *β, α* and *η* may reverse two well-known patterns of variation in the loss of phylogenetic diversity (Nee and May 1997). First, the increase in PD loss with tree imbalance can be hampered by *η* values deviating from one (Fig. 7 and Appendix 7 available as Supplementary Material). In particular when *η* < 1, this pattern results from the preferential extinction of phylogenetically redundant species in more unbalanced trees when extinction risks are clustered in large clades. Second, when *η* > 1 and *α* < 0 the loss of phylogenetic diversity proceeds faster than that of species diversity (turning their relationship from convex to concave, except in very balanced or very unbalanced trees; Fig. 6.C,E). This pattern is caused by the preferential extinction in small subtrees containing evolutionary distinct species. The only other cases where such high loss of PD is reached is when *β* < –1 and *η* > 1. This led us to introduce the notion of ‘thin ice zone’, as the region of parameters (*β* < –1 or *α* < 0; *η* > 1) for which phylogenies are prone to a sudden collapse of PD.

### Loss of phylogenetic diversity in bird family phylogenies

Our inference study shows that the phylogeny of bird families tend to exhibit *β* values comprised between –1 and 0. A similar result was found in many macroevolutionary studies, commonly observing values of *β* clustering around –1 in real phylogenies (*e.g*., Guyer and Slowinski 1991; Heard 1992; Guyer and Slowinski 1993; Slowinski and Guyer 1993; Mooers 1995; Purvis 1996; Mooers and Heard 1997; Blum and François 2006). With these topologies, we expect both *α* and *η* to play a major role in determining the potential losses of phylogenetic diversity (Fig. 6.E-F).

We observed *α* values clustering around zero, consistently with several empirical studies that found no positive relation between clade depth and clade size (Ricklefs 2007b, 2009, Rabosky et al. 2012; but see McPeek and Brown 2007). These values contrast with the value of 1 expected in Yule trees, and make phylogenies very sensitive to PD loss.

Our estimates of *η* values, based on the distribution of range sizes in bird family phylogenies, all fall between 1 and 1.5. This indicates that species in small clades tend to have smaller ranges than species in bigger clades. Range size has been shown to be one of the most important correlates of extinction risks and is one of the IUCN red list criteria (Purvis et al. 2000b; Cardillo et al. 2006; Lee and Jetz 2011; IUCN 2012; Arbetman et al. 2017).

Considering the three parameters together, we find that bird family trees are situated close to the region of the parameter space termed ‘thin ice zone’, for which we find the loss of phylogenetic diversity to be at least as fast as the loss of species diversity. In particular, the combination of negative *α* values with *η* > 1 leads to higher extinction risks for evolutionary distinct species. We can expect such a pattern as a result from evolutionary mechanisms acting simultaneously on different features of trees. For example, subtree-specific susceptibility to extinction, or stabilizing selection generating relictual lineages, are both expected to beget small subtrees with high divergence times also endowed with high species extinction risks. This pattern has been already found for past extinctions in birds using a species level measure of evolutionary distinctiveness; the authors observed in that case a similar loss of species and phylogenetic diversity (von Euler 2001; Szabo et al. 2012). Evolutionary distinct bird lineages were also shown to be more threatened by agricultural expansion and intensification than more recent lineages in Costa Rica (Frishkoff et al. 2014). This was also found in other taxa, such as marsupial mammals (Johnson et al. 2002) and *Sebastes* (Magnuson-Ford et al. 2009).

A striking result of our inference study relates to the narrow range of *α* values obtained as soon as the trees are large enough for the inference to be accurate (see the online Appendix 9 available as Supplementary Material for inferred parameter values as a function of the tip number in the phylogenies). This value, which differs from what is found in birth-death models, adds a new a new puzzle concerning the shape of empirical trees.

### Branch lengths in empirical phylogenies

The parameter *α* of the model shapes the order in which speciations take place, but does not instantiate the actual times between two consecutive speciation events, *i.e*., edge lengths. In the numerical investigations of PD loss, we considered two models for edge lengths: the pure-birth process (Yule 1925), and the Kingman coalescent (Kingman 1982). Using either of these models did not affect our results qualitatively, but affected them quantitatively (compare Fig. 4 and 6 to Figures provided in Appendices 4 and 6, available as Supplementary Material). Our modeling framework allows easy exploration of predictions under different models of edge lengths. This is interesting as many empirical phylogenies are not time-calibrated, or imprecisely. Besides, empirical phylogenetic trees were shown to often exhibit a decrease in branching tempo, *i.e*., in the rate of lineage accumulation through time (characterized in particular by estimates of the statistic 7 < 0; *e.g*., Nee et al. 1992; Zink and Slowinski 1995; Lovette and Bermingham 1999; Pybus and Harvey 2000; Rüber and Zardoya 2005; Kozak et al. 2006; Seehausen 2006; Weir 2006; McPeek 2008; Phillimore and Price 2008; Rabosky and Lovette 2008; Jønsson et al. 2012). Hence, quantitative predictions on the loss of phylogenetic diversity in the face of species extinctions could be further increased by accounting for real branch lengths. Moreover, several theoretical studies suggested that the branching tempo of species trees may change with clade age, decreasing in particular in younger clades (the ‘out of equilibrium’ hypothesis, proposed to explain the negative values of *γ* often observed in real phylogenies; Liow et al. 2010; Gascuel et al. 2015; Manceau et al. 2015; Missa et al. 2016; Bonnet-Lebrun et al. 2017). Taking into account such correlations between the age of clades and their branching tempo would also affect the expected loss of phylogenetic diversity.

The EDGE program (‘Evolutionary Distinct and Globally Endangered’; Isaac et al. 2007) encourages conservation priorities aiming at preserving most evolutionary history within the Tree of Life, by proposing a ranking of species based on combined criteria of evolutionary distinctiveness and extinction risk. Although our approach is not species-based but clade-based, it also investigates the preservation of evolutionary history based on principles linked to species evolutionary distinctiveness (related to the depths of subtrees, which depend on *α*) and to the distribution of extinction risks in the tree (which depends on *η*). Accordingly, to conserve most evolutionary history and evolutionary potential for further diversification and/or survival, priority could be given to clades that would undergo higher loss of phylogenetic diversity in the face of species extinctions, *i.e*., clades in the thin ice zone (*η* > 1 and either *β* < –1 or *α* < 0), and although not shown but only discussed herein, with *γ* < 0 (decreasing branching tempo; Pybus and Harvey 2000).

### Beyond losses of phylogenetic diversity

As we have seen earlier, the parameter *η* induces a sampling distribution on contemporary species, each species being drawn according to its frequency. In particular, our results could be interpreted in the light of rarefaction experiments (Nipperess and Matsen 2013), which study the way phylogenetic patterns in a metacommunity change as sampling decreases. Previous studies already pointed out strong impacts of non-random taxon sampling on the macroevolutionary patterns that we observe (*e.g*., Cusimano and Renner 2010). Our results provide insights on the effects of non-random sampling on phylogenetic diversity and phylogenetic tree topology. They reveal how, when the rarer species are not known, the divergence between observed and real phylogenetic diversity depends on the ranked shape of species trees, and on the relationship between relative abundances and richness of clades (being larger in particular in the thin ice zone; Fig. 6.E); and how the divergence between observed and real tree shape depends on *η* (real trees being more imbalanced if *η* < 1, and diverging from Yule trees towards more balance or more imbalance if *η* > 1; Fig. 8). These effects of incomplete sampling on macroevolutionary patterns should be particularly important to understand biodiversty patterns in bacterial and archeal phyla, which remain poorly known in particular because they likely harbor rare species having high chances to remain unnoticed.

### Conclusion

This new stochastic model of phylogenetic trees spans a large range of binary trees endowed with node rankings and species abundances/range sizes/extinction risks, based on three parameters only and interpolating other well-known one-parameter models. We showed that ranked tree shapes, non-random extinctions and the interactions thereof, may have a strong impact on the loss of phylogenetic diversity in the face of species extinctions, potentially reversing some expected patterns of variation in phylogenetic diversity. The simplicity of the model allows one to infer the parameters on empirical phylogenies. Applying our inference procedure on bird family phylogenies we found that, in this dataset, the parameters fall within a narrow range of the parameter space; and that the inferred values make the phylogenetic diversity of these trees very sensitive to species extinctions.

## Acknowledgments

The authors are very grateful to Mike Steel and Ana S.L. Rodrigues for their feedbacks on an earlier version of this manuscript. They also wish to warmly thank Arne Mooers for discussions and Walter Jetz for providing the data on bird range sizes. They thank the *Center for Interdisciplinary Research in Biology* (Collège de France, CNRS) for funding.

